# Motor sequence learning deficits in idiopathic Parkinson’s disease are associated with increased substantia nigra activity

**DOI:** 10.1101/2020.11.19.386193

**Authors:** Elinor Tzvi, Richard Bey, Matthias Nitschke, Norbert Brüggemann, Joseph Classen, Thomas F. Münte, Ulrike M. Krämer, Jost-Julian Rumpf

## Abstract

Previous studies have shown that persons with Parkinson’s disease (pwPD) share specific deficits in learning new sequential movements, but the neural substrates of this impairment remain unclear. In addition, the degree to which striatal dopaminergic denervation in PD affects the cortico-striato-cerebellar motor learning network remains unknown. We aimed to answer these questions using fMRI in 16 pwPD and 16 healthy age-matched control subjects while they performed an implicit motor sequence learning task. While learning was absent in both pwPD and controls assessed with reaction time differences between sequential and random trials, larger error-rates during the latter suggest that at least some of the complex sequence was encoded. Moreover, we found that while healthy controls could improve general task performance indexed by decreased reaction times across both sequence and random blocks, pwPD could not, suggesting disease-specific deficits in learning of stimulus-response associations. Using fMRI, we found that this effect in pwPD was correlated with decreased activity in the hippocampus over time. Importantly, activity in the substantia nigra (SN) and adjacent bilateral midbrain was specifically increased during sequence learning in pwPD compared to healthy controls, and significantly correlated with sequence-specific learning deficits. As increased SN activity was also associated (on trend) with higher doses of dopaminergic medication as well as disease duration, the results suggest that learning deficits in PD are associated with disease progression, indexing an increased drive to recruit dopaminergic neurons in the SN, however unsuccessfully. Finally, we found no differences between pwPD and controls in task modulation of the cortico-striato-cerebellar network. Notably, in both groups Bayesian model selection revealed cortico-cerebellar connections modulated by the task, suggesting that despite behavioral and activation differences, the same cortico-cerebellar circuitry is recruited for implementing the motor task.

## 1. Introduction

The clinical diagnosis of Parkinson’s disease (PD), the second most prevalent neurodegenerative disorder worldwide, is based on the cardinal motor symptoms bradykinesia, muscle rigidity, and tremor (Berardelli et al., 2013). These PD-defining motor symptoms are mainly related to the degeneration of dopaminergic neurons in the substantia nigra (SN) pars compacta (Fearnley and Lees, 1991) and loss of neurons in the nigro-striatal pathway projecting primarily to the putamen (Dauer and Przedborski, 2003). However, PD also affects several other neurotransmitter systems, including cholinergic, noradrenergic, glutaminergic, and GABAergic pathways (Brichta et al., 2013; Calabresi et al., 2006; Francis and Perry, 2007). Persons with PD (pwPD) often also suffer from cognitive dysfunctions that compromise working memory, executive functions and attention (Williams-Gray et al., 2007) as well as implicit motor learning and retention of motor skills (Redgrave et al., 2010). Early procedural learning studies showed that pwPD have specific learning-related deficits (Doyon et al., 1997; Jackson et al., 1995; Pascual-Leone et al., 1993). These deficits were already evident in very early stages of the disease and even detectable when the asymptomatic hand was tested in unilateral de novo pwPD (Dan et al., 2015). However, these findings were not always consistent (Seidler et al., 2007; Smith et al., 2001; Werheid et al., 2003), probably due to strong variability in disease progression (Muslimović et al., 2007; Stephan et al., 2011) and pharmacotherapy (Kwak et al., 2012, 2010) as well as variability in task demands. A meta-analysis revealed that while pwPD have clear deficits in their ability to implicitly learn a motor sequence compared to healthy controls, they were still able to improve their performance with practice, albeit to a lesser degree (Hayes et al., 2015).

To investigate the underlying neural mechanisms of motor sequence learning (MSL) deficits in PD, previous research has employed imaging methods such as positron emission tomography and functional magnetic resonance imaging (fMRI). Early studies showed that acquisition and retention of a motor sequence recruits different brain regions in PD compared to controls (Mentis et al., 2003; Nakamura et al., 2000) and leads to over-activation of premotor and parietal areas as well as cerebellum in PD (Caproni et al., 2013; Sabatini et al., 2000; Wu and Hallett, 2005). Effective connectivity analysis showed that interactions between cerebellar and premotor areas were significantly reduced in PD during the automatic phase of MSL (Wu et al., 2010). Thus, evidence suggests that increased activity and decreased connectivity within the motor network may underlie MSL deficits in PD.

In healthy subjects, learning a motor sequence activates a distributed network including striatum, thalamus as well as motor cortical areas, parietal cortex, dorsolateral prefrontal cortex and cerebellum (for review see: (Hardwick et al., 2013)). Theoretical models suggest that motor learning is implemented through specific cortico-striatal and cortico-cerebellar circuits, which mediate different learning stages (Doyon et al., 2009). Previously, we tested this model in healthy subjects and found that learning negatively modulated connections from M1 to cerebellum (Tzvi et al. 2014) but also between putamen and cerebellum (Tzvi et al. 2015; Tzvi et al. 2017). This suggests that besides the cerebellum, the putamen may also play an important role in acquisition of new motor sequences. In this study, we aimed to investigate how the dysfunctional nigro-striatal dopaminergic system in PD affects neural activity and connectivity in a motor learning network, while pwPD perform an implicit MSL task, concurrent to fMRI. We hypothesized that dysfunction of the nigro-striatal system in PD will affect activity and connectivity in the cortico-striato-cerebellar network. Specifically, we expected that learning deficits in pwPD would be associated with neural changes in the striatum, and lead to altered network interactions underlying MSL.

## 2. Materials and methods

### 2.1. Participants

Twenty-three persons with Parkinson’s disease (see Table 1 for demographics) volunteered to participate in the study. Idiopathic PD was diagnosed by expert neurologists (M.N. & N.B.) and the severity of motor symptoms was assessed according to the Unified Parkinson’s disease Rating Scale part III (UPDRS III; (Fahn et al., 1987). PwPD were recruited from the outpatient clinic of the Department of Neurology of the University Hospital of Lübeck. Upon recruitment, pwPD were first tested for their general cognitive abilities using the Mini-Mental State Examination (Pangman et al., 2000). Those who scored more than 20 out of 30 points on the Mini-Mental State Examination and less than 40 points on UPDRS III were eligible to participate. In addition, we used a short pre-task test block to ensure that finger-tapping skills required to perform the task were intact. From this cohort, we had to exclude five pwPD due to diagnosis of depression, or severe cortical atrophy detected in the anatomical scan (see Table 1 for more details). Two additional pwPD were rejected from analysis of task-fMRI data due to bad performance (one fell asleep during the task and the other made more than 50 % errors). Thus, we included sixteen pwPD (10 males; age: 46-77; mean age: 64.3). To this sample we age-matched sixteen neurologically healthy controls (7 males; age: 53-77; mean age: 62.6) who were recruited from the general community. All participants were right-handed (except for one pwPD) and had normal or corrected to normal vision. Participants gave informed written consent prior to study participation. The study was approved by the Ethics Committee of the University of Lübeck.

**Table 1:** Characteristics of persons with PD.

| ID | age | gender | MMSE | UPDRSIII | DD | LED |
| --- | --- | --- | --- | --- | --- | --- |
| *P01 | 48 | m | 30 | 20 | 7 | 760 |
| P02 | 65 | f | 22 | 10 | - | 325 |
| P03 | 74 | m | 24 | 22 | 13 | 1169 |
| *P05 | 53 | f | 30 | 16 | 8 | - |
| P06 | 46 | m | 29 | 10 | 7 | 565 |
| P07 | 70 | f | 29 | 21 | 9 | 250 |
| P09 | 65 | m | 30 | 34 | 3 | 179 |
| P10 | 54 | m | 25 | 34 | 3 | 905 |
| **P11 | 66 | m | 25 | 32 | 5 | 782 |
| P12 | 61 | m | 28 | 36 | 5 | 814 |
| **P13 | 48 | m | 27 | 25 | 5 | 1246 |
| P15 | 69 | m | 28 | 39 | 10 | 450 |
| P16 | 56 | f | 29 | 37 | 2 | 210 |
| *P18 | 75 | m | 26 | 21 | 2 | 563 |
| P19 | 60 | m | 29 | 30 | 1 | 100 |
| P30 | 69 | m | 30 | 11 | 10 | 766 |
| P33 | 77 | f | 29 | 11 | 6 | 430 |
| P35 | 70 | m | 30 | 14 | 19 | 650 |
| P36 | 61 | f | 29 | 9 | 3 | 400 |
| P37 | 61 | f | 30 | 5 | 12 | 726 |
| P38 | 70 | m | 28 | 8 | 3 | 320 |

\* Participants excluded from the study
\*\* Participants excluded from behavioral, task-based fMRI analyses and DCM analysis
- Missing data

### 2.2. Experimental paradigm and task design

Participants performed a modified version of the serial reaction time task (Nissen and Bullemer, 1987) while lying supine in the magnetic resonance imaging (MRI) scanner after a short familiarization with the task. PwPD were under their standard dopaminergic medication (Table 1) during task performance. The visual stimuli were delivered to the participants through MR-compatible goggles. In each trial, four squares were presented in a horizontal array, with each square (from left to right) associated with the following four fingers: middle finger left hand, index finger left hand, index finger right hand, middle finger right hand (Fig. 1A).

**Figure 1.**
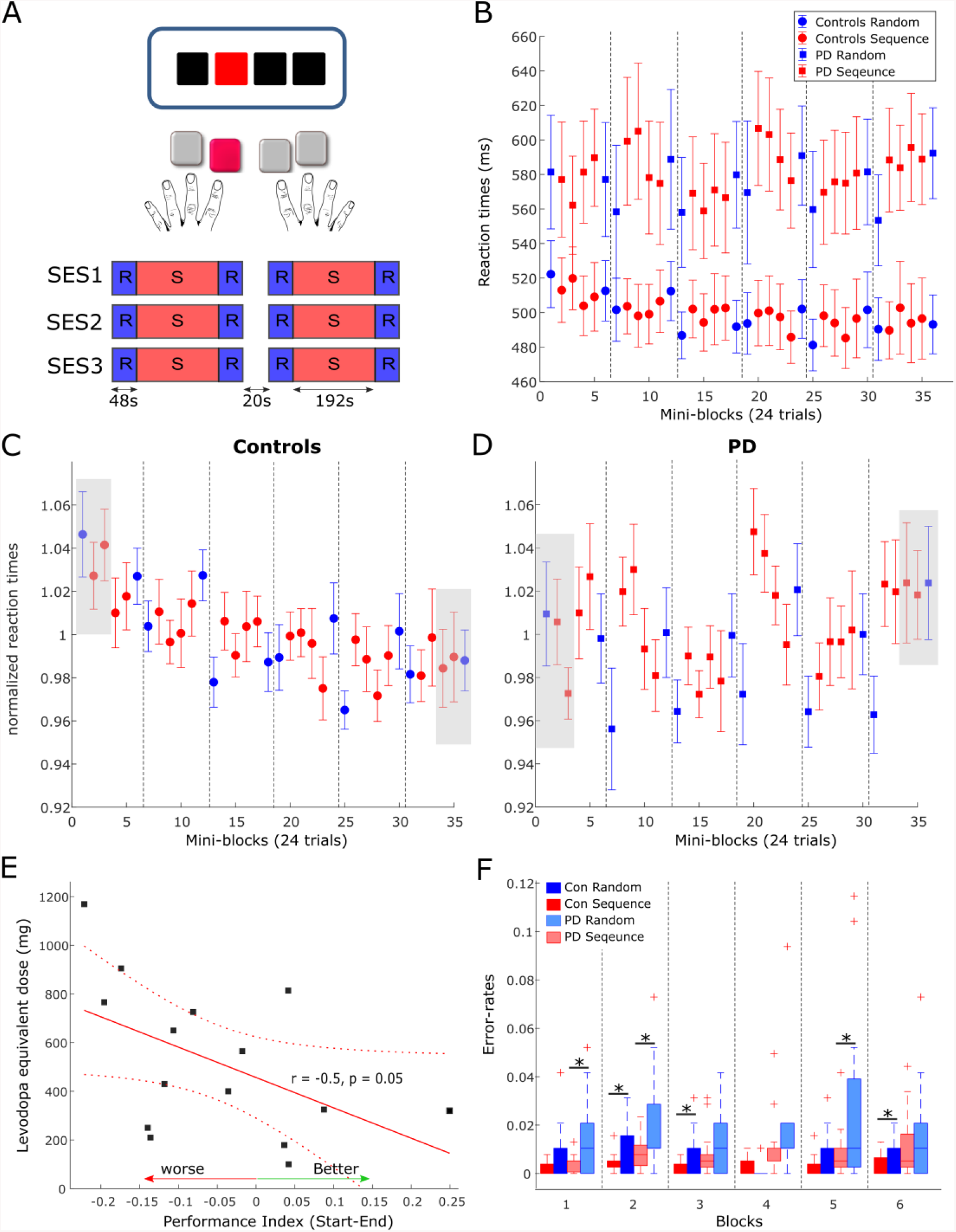
**A** Task and experimental design. In each trial, subjects viewed three black squares and one red square which served as the target. Subjects were instructed to press the button corresponding to the location of the red square. The task consisted of three MRI sessions (SES1-SES3) including two blocks each, amounting to a total of six blocks. 20 sec breaks were introduced between sessions and blocks. **B** Reactions times in PD (squares) and controls (circles) in each condition and each mini-block of 24 trials. Error-bars are standard error of the mean across subjects in each group. Dashed grey lines depicts the breaks between the blocks. **C-D** Normalized reaction times in healthy controls (C) and PD (D). Grey boxes show the blocks used for comparison of performance between groups. **E** Correlation between the performance index, calculated as the differences between the first three blocks of normalized RTs and the last three blocks, and the levodopa equivalent dose. **F** Error rates in each condition and each block for PD (light colors) and healthy controls. Asterisks mark significant differences within the group.

Participants were instructed to respond to the red coloured square with the corresponding button on an MRI-compatible keypad, one for each hand, as precisely and quickly as possible. Stimuli were presented either in a pseudorandom order (RND) or as a 12-items-sequence (SEQ, “1-2-1-4-2-3-4-1-3-2-4-3”). Participants were not aware of the existence of any patterns in the stimuli. Pseudorandom orders were generated using Matlab (Natick, MA) such that items were not repeated. The task consisted of three MRI sessions including two blocks each, amounting to a total of six blocks (Fig. 1A). Each block contained eight repetitions of the 12-element sequence (i.e. 96 trials) as well as 24 random trials before and after the sequence block. A 20 sec break was introduced between the blocks during which participants were instructed to fixate on a black cross in the center of the goggle screen. Visual stimuli were presented until the onset of a button press or the onset of the next trial. The inter-stimulus interval was 2000 ms. We used Presentation^®^ software (Version 16.3, www.neurobs.com) to present stimuli and to synchronize the stimulus presentation and the MR functional sequences.

### 2.3. MRI data acquisition and pre-processing

The MR data were recorded using a 3T Philips Achieva head-scanner at the Institute of Neuroradiology, University of Lübeck. Functional MRI (fMRI) data (T_2_*) were collected using blood oxygen level dependent (BOLD) contrast in three sessions with 300 volumes each and a gradient-echo EPI sequence following these specifications: repetition time TR = 2000 ms, echo time TE = 30 ms, flip angle = 90°, matrix size 64 × 64, FOV = 192 × 192 mm with a whole brain coverage, ascending slice order of 3 mm thickness, and a 0.75 mm gap to avoid crosstalk between slices, in-plane resolution of 3 × 3 mm, and SENSE factor of 2. Subsequently, a high resolution T_1_-weighted structural image was acquired with FOV = 240 × 240 mm; matrix = 240 × 240; 180 sagittal slices of 1 mm thickness.

Preprocessing of fMRI data was performed using SPM12 software package (http://www.fil.ion.ucl.ac.uk/spm/). The preprocessing included correction for differences in image acquisition time between slices, a six-parameter rigid body spatial transformation to correct for head motion during data acquisition, co-registration of the structural image to the mean functional image, grey and white matter segmentation, and spatial normalization of the structural image to a standard template (Montreal Neurological Institute, MNI). In order to reduce the influence of motion and unspecific physiological effects, a regression of nuisance variables from the data was performed. Nuisance variables included white matter and ventricular signals and the six motion parameters determined in the realignment procedure. Spatial normalization of the functional images was applied using the normalization parameters estimated in the previous preprocessing step and resampling to 3 × 3 × 3 mm. Finally, spatial smoothing with a Gaussian kernel of 8 mm full width half maximum was applied.

### 2.4. Task-based fMRI statistical analysis

Imaging data was subsequently modeled using the general linear model (GLM) in a block design manner. Linear regressors were obtained for each of the experimental conditions (SEQ and RND) and each session (SES1, SES2, SES3) in each participant. First level GLM analysis thus contained six experimental blocks, two for each session, modelled as a box function with the duration of each block and convolved with a hemodynamic response function. Movement related parameters from the realignment process were included in the GLM as regressors of no-interest to account for variance caused by head motion. We applied a high-pass filter (256 sec) to remove low-frequency noise. First-level contrast images were generated using a one-sample t-test against rest periods between the blocks.

We analyzed contrast images from each participant on the second level using a random effects model. To investigate general task-related differences in activity between the groups, we performed a flexible factorial analysis accounting for main effects of Group (PD, Controls), Session (SES1, SES3), and Group x Session interaction. In addition, we explored group differences in sequence-learning activity by collapsing data across all SEQ sessions and all RND sessions and performing a flexible factorial analysis accounting for main effects of Group (PD, Controls), Condition (SEQ, RND), and interaction effects. Statistical significance was established using a whole-brain voxel-level threshold of p = 0.001 and cluster level of p < 0.05, FWE corrected over the entire cluster. To analyze interactions effects, we used rfxplot toolbox (Gläscher, 2009) to extract the contrast estimates from significant voxels in a sphere (radius = 4 mm) around the peak activity.

We additionally calculated a metric for head motion using the translation parameters: × – left/right, y – anterior/posterior, z – superior/inferior. These parameters represent frame-wise displacement in 3D: 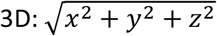. This displacement parameter was then averaged across all volumes in each session and then across sessions to produce a mean motion parameter in mm for each subject. We then compared the mean motion parameter between pwPD and controls using a two-sample t-test. One control subject was excluded from the fMRI analysis due to excessive head movements resulting in a sample of 15 healthy controls and 16 pwPD. Note that this subject was included in the behavioral analyses. After this exclusion, analysis of head motion using displacement revealed no differences (p > 0.1) between PD (0.69 ± 0.41) and controls (0.67 ± 0.44).

### 2.5. Dynamic causal modelling

We used dynamic causal modeling (DCM, Friston et al., 2003) as implemented in SPM12 (v. 7771), version DCM12.5, to investigate changes in effective connectivity within a cortico-striato-thalamo-cerebellar network due to impaired learning and motor performance in PD. Importantly, we hypothesized that connectivity patterns within this network would differ in PD compared to healthy controls. To this end, 12 nodes were specified containing bilateral motor cortical areas: primary motor cortex (M1), supplementary motor area (SMA) and premotor cortex (PMC) as well as putamen, thalamus and cerebellum. Within each hemisphere and the contralateral cerebellum, all VOIs were assumed to be fully connected (Fig. 2A). We also allowed an intrinsic connection between homolog M1 based on previous work (Tzvi et al., 2017). In addition, our previous fMRI studies of MSL showed that specifying the right cerebellum as input node leads to highest exceedance probability (Tzvi et al., 2017, 2015, 2014). We therefore chose this node as input here as well. Note that it is likely that the driving input is mediated by un-modelled regions such as the visual cortex. The input and connections specified here do not necessarily represent anatomical input and connectivity but rather a “net effect”. We then specified 14 different models which allowed modulation of different connections (Fig. 2B). These were based on:

**Figure 2.**
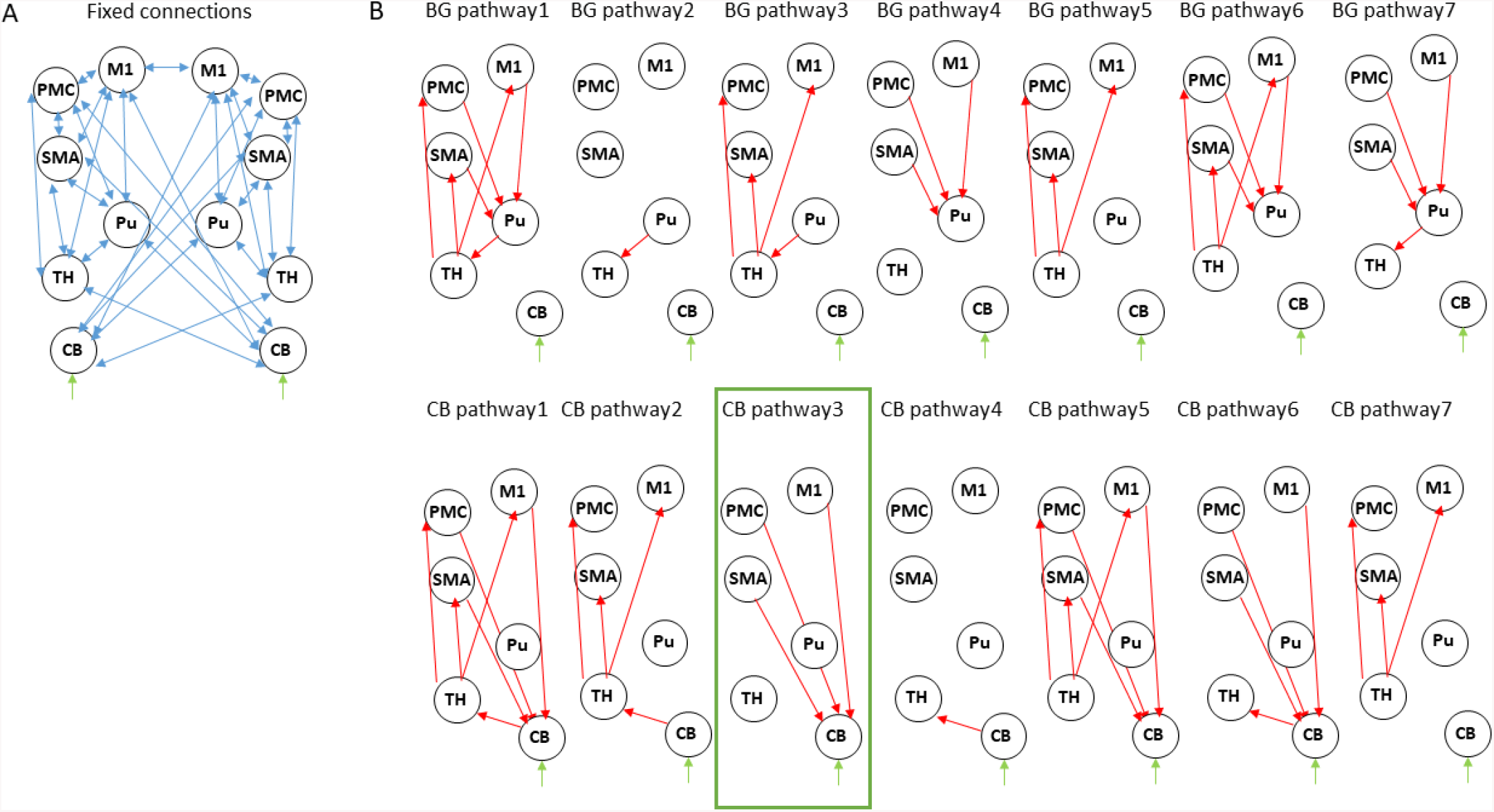
Dynamic causal modelling. **A** Intrinsic connections across all models. **B** Modulatory connections. For simplicity we show only one hemisphere but the modulatory connections are assumed in both. The upper row describes models within the "basal ganglia pathway” with different modulations of the motor-putamen-thalamo-motor circuit. The bottom row describes models within the "cerebellar pathway” with different modulations of the motor-cerebello-thalamo-motor network. M1-primary motor cortex, Pu – putamen, SMA – supplementary motor area, PMC – premotor cortex, TH – thalamus, CB – cerebellum. The winning model in both pwPD and healthy controls is marked with a green frame.

1. “Basal-ganglia pathway”: these models are based on the classical model of direct and indirect pathways in the basal ganglia underlying motor actions. According to this model, cortical activation leads to activation of striatum, which in turn leads to disinhibition of the thalamus which projects to the cortex (Calabresi et al., 2014). The seven models specified here model modulation by the motor task of the entire circuit (model BG pathway 1) or of parts of the circuit (model BG pathway 2-7). In PD, dopaminergic denervation in SN leads to abnormal activation of striatal output nuclei and thus an over-inhibition of thalamic neurons projecting to the motor cortex.
2. “Cerebellar pathway”: here, we specified a network based on the cerebello-thalamo-cortical pathway connecting sensorimotor and premotor areas to cerebellum, from cerebellum to thalamus and from thalamus back to sensorimotor and premotor areas (Ramnani, 2006). Here as well, we modelled modulation of the entire circuit (model CB pathway 1) or of parts of the circuit (model CB pathway 2-7).

For each model, a one-state bilinear system of differential equations was inverted and together with a biophysically motivated hemodynamic model, an estimated BOLD signal was produced. This modelled BOLD signal was then iteratively fitted to the real data through a gradient ascent on the free-energy bound. We describe the results of the DCM analysis on two levels: on a model level and on a parametric level. First, we selected a “winning” model out of a candidate set of equally plausible models, based on its protected exceeding probability, using Bayesian model selection. This was done for each group separately. Second, connectivity parameters within the “winning model” were then analyzed using a 2 × 2 mixed effects ANOVA with factors COND (SEQ, RND) and group (PD, controls) to discover learning-specific changes due to PD. We extracted the modulatory parameters from the winning model by averaging across the three sessions. Note that in some cases, the model did not converge, leading to zero parameter estimates. In order to not bias the results towards zero, these sessions were identified using the spm_dcm_fmri_check.m script and removed from further analyses. In one pwPD and one healthy control, the model did not converge in any session, leading to exclusion of these two subjects for the DCM analyses only.

#### 2.1.1. Time series extraction

We specified the following as volumes of interest (VOIs): primary motor cortex (M1), supplementary motor area (SMA), premotor cortex (PMC), putamen (Pu), Thalamus (TH) and cerebellum (CB). Time series were extracted across all experimental blocks and sessions, in order to account for both learning- and non-learning related changes in the BOLD signal. The coordinates of the sphere centers for each VOI were selected based on the local maxima of the group level task vs. baseline contrast (see Table 2). Using a singular value decomposition procedure implemented in SPM12, we computed the first eigenvariate across voxels within 4 mm radius from the sphere center for each subject. Time series were then detrended and sharp improbable temporal artifacts were smoothed by an iterative procedure implementing a 6-point cubic-spline interpolation. Finally, we estimated the explained variance of the signals we extracted from each of the VOIs by computing the proportion of the first eigenvariate in the signal. The minimal variance explained (across all subjects and VOIs) was 62% which means that the first eigenvariate explained much of the signal variance in the different VOIs.

**Table 2:** Dynamic causal modelling nodes: Task > Baseline.

| <u>Region</u> | <u>coordinates</u> | <u>Sub-region</u> | <u>t-val</u> |
| --- | --- | --- | --- |
| L M1 | -54 -16 32 | BA4 | 7.03 |
| R M1 | 56 -16 40 | BA4 | 4.65 |
| L SMA | -6 4 56 | BA6 | 9.43 |
| R SMA | 0 2 48 |  | 9.85 |
| L PMC | -42 -8 60 | BA6 | 7.68 |
| R PMC | 48 -4 56 | BA6 | 8.34 |
| L Putamen | -24 -2 6 |  | 6.87 |
| R putamen | 26 -2 6 |  | 6.59 |
| L cerebellum | -24 -66 -56 | lobule VIII | 4.65 |
| R cerebellum | 16 -64 -54 | lobule VIII | 4.16 |
| L Thalamus | -14 -16 8 |  | 4.08 |
| R Thalamus | 14 -14 6 |  | 2.39 |

## 3. Results

### 3.1. Behavioral results

#### Reaction times

We analyzed changes in reaction times (RT) due to learning and PD using a repeated-measures analysis of variance (rmANOVA) with factors Group (PD, controls), Condition (SEQ, RND) and Block (1-6). Generally, pwPD were significantly slower in responding to the stimulus, across both SEQ and RND blocks, compared to healthy controls, as reflected by a main effect of Group (F_1,30_ = 5.5, p = 0.03, Fig. 1B). No other main effects or interactions were found (p > 0.1). We then investigated whether subjects of both groups learned the underlying motor sequence using two separate rmANOVA with factors Condition and Block in each group. Neither PD nor controls showed specific sequence learning effects reflected by either a main effect of Condition (both p > 0.1) or a significant Condition x Block interaction (both p > 0.2).

However, a main effect of Block in the control group (F_5,75_ = 2.8; p = 0.02) suggested general task improvement independent of the condition (Fig. 1C), which was not evident in PD (p = 0.8, Fig. 1D). We then used a two-sample t-test to explore whether task improvement differed between PD and controls. First, RT were normalized by the averaged RT across the task in each subject (Fig. 1C-D). Changes in normalized RT across the task thus represent general learning effects, irrespective of individual RT performance. Then, we specified a Performance Index using the difference between normalized RTs averaged across the first three blocks of the task (one random and two sequence blocks with 24 trials each) and normalized RT averaged across the three last blocks of the task (see grey squares in Fig. 1C-D). Indeed, we found that the Performance Index was larger in controls compared to PD (t_30_ = 2.1, p = 0.04), suggesting the pwPD had deficits in learning stimulus-response associations in this task. This effect was not related to motor deficits in PD, evaluated using the UPDRS III score (p > 0.8) but did tend to relate to levodopa equivalent dose (LED) such that larger LED was associated with worse performance at the end of the task (Fig. 1E, r = -0.48, p = 0.057). There was no correlation between the Performance Index and disease duration (p > 0.2).

#### Error rates

In terms of error-rates, pwPD produced significantly more errors compared to healthy controls (Z = 2.75, p = 0.006; Fig. 1F). Similar to the RT analysis above, condition differences in error-rates were assessed in each group separately. PwPD produced significantly more errors (corrected for multiple comparisons) in RND compared to SEQ in block 1 (Z = 2.59, p < 0.01), block 2 (Z = 2.36, p = 0.02), and block 5 (Z = 2.71, p = 0.007). In healthy controls, error rates also tended to be larger in RND compared to SEQ in block 2 (Z = 2.30, p = 0.02), block 3 (Z = 1.96, p < 0.05), and block 6 (Z = 2.59, p < 0.01) but this difference did not survive correction for multiple comparisons. These differences indirectly suggest that some learning of the sequence has taken place in both groups.

In sum, while RT analysis showed no condition differences in both healthy controls and pwPD, the significantly higher error-rates during RND blocks indicate that some learning of the sequence has taken place. Healthy controls improved condition unspecific task performance as indicated by faster RTs at the end of the task compared to the beginning whereas pwPD could not. This suggests that pwPD are impaired in learning stimulus-response associations.

### 3.2. Functional MRI results

#### Decrease in left hippocampus activity associated with general task-related deficits in PD

To investigate changes in general task-related activity due to PD, we subjected the contrasts of Session 1 and Session 3 in each group to a flexible factorial design with factors Group (PD, controls) and Session (SES1, SES3). There were no significant clusters for the Group factor. Although no significant Group x Session interactions were observed on a family-wise error (FWE) corrected p-level, a few large clusters in right premotor cortex, left insula, left cerebellum crus II, and left hippocampus were evident on an uncorrected p < 0.001 voxel-based significance level (see Table 3 and Fig. 3A). Group × Session interactions show regions associated with both task-related changes over time as well as differences between PD and controls (Fig. 3B). Post-hoc paired t-tests show that these interactions arise from a specific increase from SES1 to SES3 in healthy controls (Fig. 3B, left cerebellar crus II: t_14_ = 3.9, p = 0.002; right premotor cortex: t_29_ = 3.4, p = 0.004; left hippocampus: t_29_ = 2.2, p = 0.04), with the opposite pattern in PD (Fig. 3B, right premotor cortex: t_15_ = 2.5, p = 0.02, left hippocampus: t_15_ = 2.6, p = 0.02, left cerebellar crus II: ns). In addition, two-sample t-tests comparing both groups revealed decreased activity in PD compared to controls in SES3 (Fig. 3B, left cerebellar crus II: t_29_ = 2.8, p = 0.008; right premotor cortex: t_29_ = 3.1, p = 0.004; left hippocampus: t_29_ = 2.9, p = 0.007). These results suggest that healthy controls recruit regions important for task performance, whereas in pwPD, decreased activity in these regions could be related to deficits in general task performance.

**Table 3:** fMRI task activations (voxel level: p < 0.001)

| Region | MNI-coordinates |  |  | t-value | n-voxels |
| --- | --- | --- | --- | --- | --- |
| **Group (PC, Cont) x SES (1,3)** |  |  |  |  |  |
| Right premotor cortex | 50 | -4 | 54 | 4.33 | 197 |
| Right inferior frontal gyrus | 34 | 8 | -12 | 4.23 | 25 |
| Left superior temporal gyrus | -42 | -24 | -4 | 4.17 | 63 |
| Left cerebellum crus II | -32 | -76 | -46 | 4.12 | 100 |
| Left Insula | -36 | 12 | -14 | 4.04 | 110 |
| Right superior temporal pole | 54 | 12 | -20 | 3.85 | 12 |
| Left hippocampus | -26 | -16 | -10 | 3.78 | 89 |
| **Group (PC, Cont) x Cond (SEQ, RND)** |  |  |  |  |  |
| *Right substantia nigra | 14 | -18 | -6 | 4.47 | 12 |
| *Right brainstem | 8 | -30 | -4 | 4.02 | 205 |
| *Left brainstem | -6 | -32 | -6 | 3.91 | 162 |
| Left superior parietal lobule | -20 | -52 | 70 | 4.07 | 65 |
| Right hippocampus | 24 | -22 | -14 | 3.75 | 23 |
| Left thalamus | -12 | -12 | 0 | 3.45 | 16 |

\* $p = 0.05$ , family-wise error corrected across the cluster

**Figure 3.**
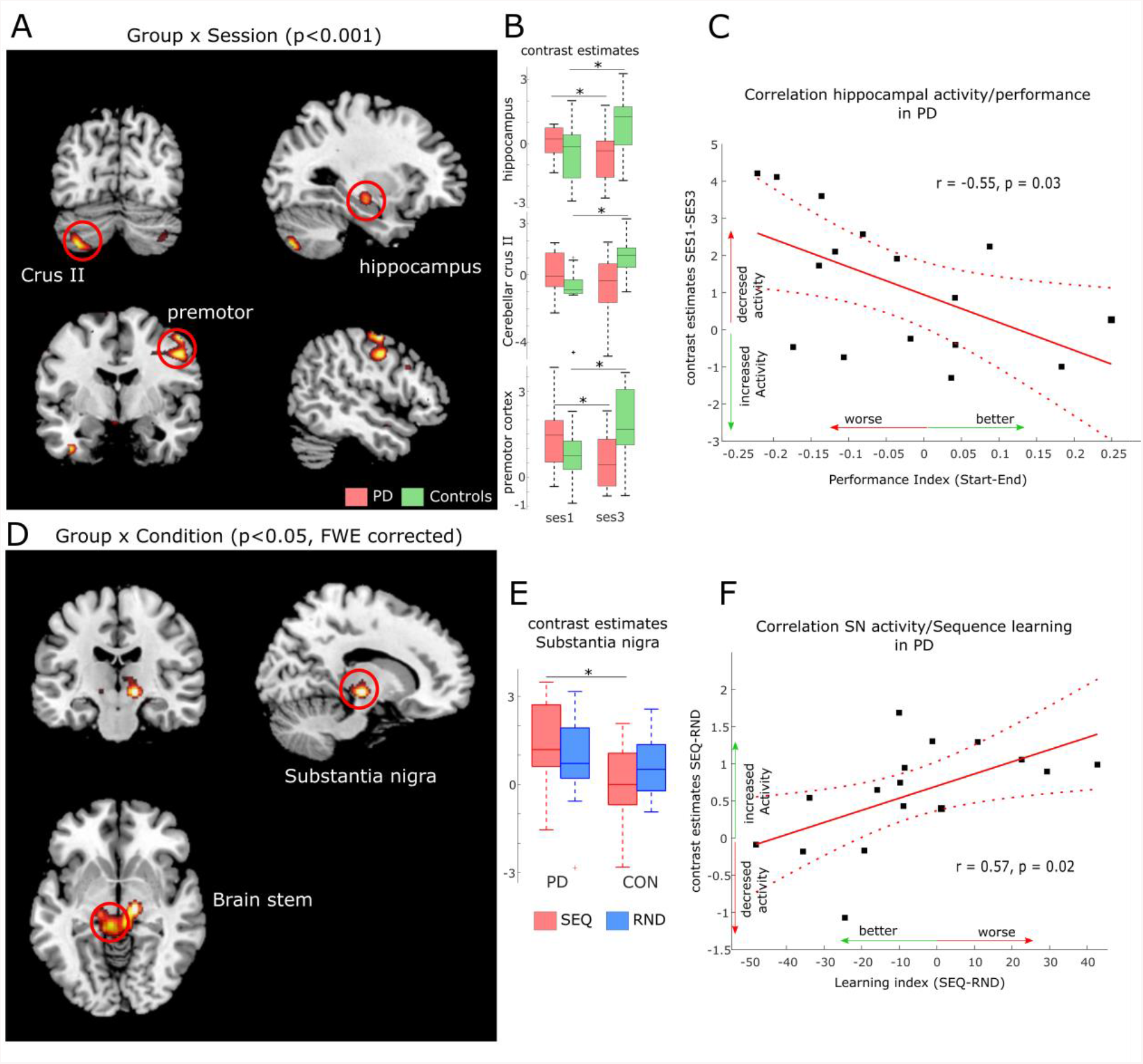
FMRI results. **A** Activation maps for Group (PD, controls) x Session (SES1, SES3) interactions at p < 0.001, uncorrected voxel-level threshold. **B** Contrast estimates derived from activation clusters shown in A. Boxplots show significant increased activity in controls and decreased activity in PD. **C** Correlation between activity increase in hippocampus and improved performance in PD. **D** Activation maps for Group (PD, controls) x Condition (Sequence, Random) interactions at p < 0.05, cluster-level family-wise error corrected. **E** Contrast estimates derived from the activation cluster in substantia nigra (SN) shown in D. Boxplots show significant increased activity in PD compared to controls during sequence blocks. **F** Correlation between activity increase in SN and worse sequence learning in PD.

To test this hypothesis, we explored two relationships in the PD group. First, we investigated a link between activity changes activity from SES1 to SES3 and the Performance Index (see Table 4) and second, we explored a link between activity changes activity from SES1 to SES3 and measures of disease progression, namely LED and disease duration. We found a negative correlation between activity changes in left hippocampus and the Performance Index (Fig. 3C, r = -0.55, p = 0.03), which meant that pwPD who slowed towards the end of the task, i.e. performed worse, also showed relatively decreased left hippocampus activity in SES3 compared to SES1. Second, we found a positive correlation between activity changes in left cerebellar crus II and LED (r = 0.53, p = 0.03, supp. Fig. 1A) as well as disease duration (r = 0.59, p = 0.02, supp. Fig. 1B). This suggests that increased dopaminergic denervation, reflected in disease duration and higher daily doses of dopaminergic medication, is associated with decreased left cerebellar crus II activity from SES1 to SES3. Together, these results suggest that impaired learning of stimulus-response associations in PD is associated with a hippocampal decrease in activity, whereas the severity of the disease affects recruitment of left cerebellar crus II.

**Table 4:** behavioral measures in the serial reaction time task.

| Name | Definition |
| --- | --- |
| Performance Index | Normalized RTs in each subject. Represents individual differences in performance across time (Fig. 1C-E, Fig. 3C). Measured as the relation between RTs in each mini-block divided by the averaged RT across the entire task. |
| Learning Index | Difference in reaction-times between sequence and random blocks (Fig. 3F, Fig. 4D-E). Measured as the differences in averaged RTs across blocks 3-6, between sequence and random. |

#### Increased SN activity associated with sequence learning deficits in PD

Based on the behavioral results (analysis of error-rates) that indirectly suggest sequence-specific learning in both groups, we assessed possible differences in neural responses to implicit MSL in PD compared to controls, using a flexible factorial design with factors Group and Condition (SEQ vs. RND across all sessions). We found a Group × Condition interaction in a large cluster with peak activity in left substantia nigra (SN, t_59_ = 4.47, p < 0.05, FWE cluster-level corrected) and adjacent bilateral midbrain areas (Table 3, Fig. 3D). Post-hoc paired t-tests show that this interaction was caused by decreased activity in healthy controls compared to relatively increased activity in PD during SEQ blocks (Fig. 3E, t_29_ = 2.7, p = 0.01). Differences in RND were non-significant (p = 0.5).

To determine whether these differences in neural activity were related to individual sequence learning, we first created a Learning Index based on RT differences between SEQ and RND averaged across SES2 and SES3 (blocks 3-6). Note that this Learning Index is different than the Performance Index specified above (Table 4). We then correlated the Learning Index with learning-related activity (SEQ-RND) in SN. We found a significant positive correlation (r = 0.57, p = 0.02; Fig. 3F) in the PD group, which meant that stronger activity in sequence blocks was driven by reduced learning. In addition, learning-related (SEQ-RND) activity in SN tended to positively correlate with disease duration (r = 0.51, p = 0.05, supp. Fig. 2B) as well as levodopa equivalent dose (r = 0.47, p = 0.06, supp. Fig. 2A). This suggests that the severity of the disease, reflected in its duration and higher dopaminergic medication, leads to increased activation of SN during learning of a motor sequence. There was no association between learning-related (SEQ-RND) activity in SN and the UPDRS score (p > 0.1).

### 3.3. Dynamic causal modelling

#### Condition differences in modulation of motor and premotor areas to cerebellum

Using Dynamic Causal Modelling (DCM), we next asked whether striatal dysfunction in pwPD leads to changes in connectivity in a cortico-striato-cerebellar network underlying motor learning. To this end, we specified two pathways, along which modulation by learning and by task performance may be evident: First, a “basal ganglia circuit”, connecting motor and premotor cortex to putamen, then to thalamus and back to motor cortical regions; second, a “cerebellar circuit” connecting motor and premotor cortex to cerebellum, then to thalamus and back to motor cortical regions (see Fig. 2B). A total of 14 models were compared using Bayesian model selection, 7 models for each pathway (Fig. 2B). Random-effects Bayesian model selection showed that model 10 (Fig. 4A) had the strongest protected exceedance probability (Fig. 4C) in both PD and controls (PD: 0.50, Controls: 0.64, see comparison to other models in Fig. 4C). In this model, shown in Fig. 4A, connections from bilateral M1, premotor cortex (PMC) and supplementary motor area (SMA) to contralateral cerebellum (CB) were modulated by the motor task (both SEQ and RND blocks). Next, we tested differences in modulatory parameters between PD and controls in the six connections of the winning model (bilateral M1, SMA and PMC -> CB), using a 2 × 2 mixed effects ANOVA with factors COND (SEQ, RND) and group (PD, controls). We found in all connections (except for right M1 -> left CB), a significantly larger negative modulation in RND compared to SEQ (Fig. 4B, Table 5). There were no differences between PD and controls in terms of modulatory parameters (p > 0.1 for the group factor for all modulatory connections).

**Table 5:**
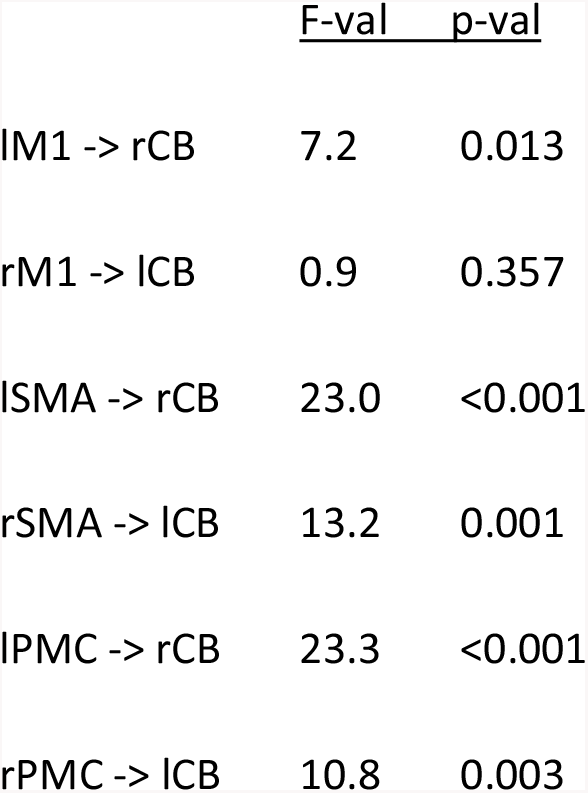
Modulatory parameters in the winning model (main effect of COND: RND > SEQ)

**Figure 4.**
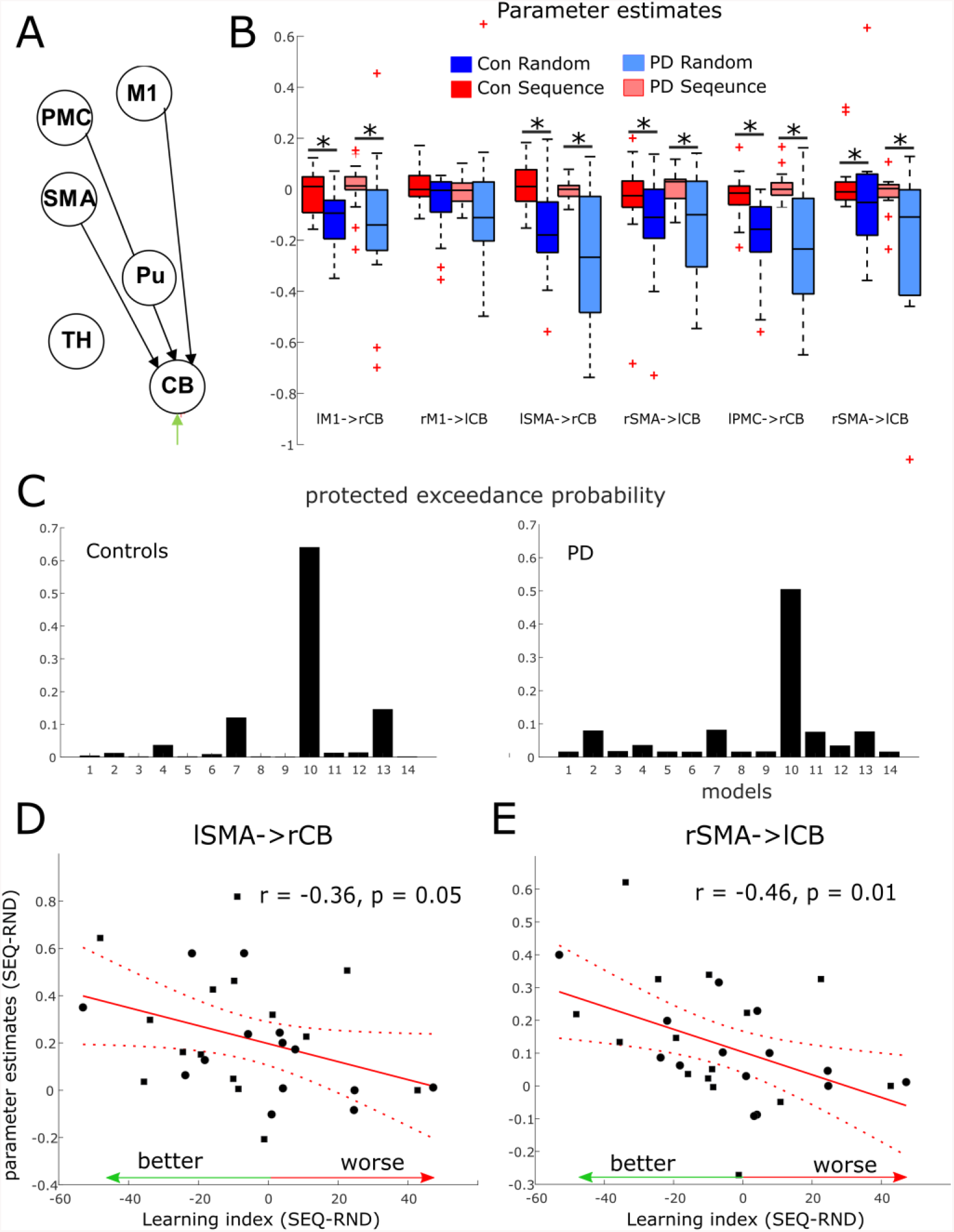
DCM results. **A** Winning model in both groups. Note that the model had 12 nodes in both hemispheres. Here only 6 nodes in left cerebrum and right cerebellum are depicted. M1 – primary motor cortex. PMC – premotor cortex. SMA – supplementary motor area. Pu – putamen. TH – thalamus. CB – cerebellum. **B** Boxplots for parameter estimates in the winning model (A) for both groups (PD, controls) and conditions (Sequence, Random). Significant differences are marked with a start. **C** Protected exceedance probability in healthy controls and PD showing model 10 (in A) was favorable over the other models (Fig. 2). **D-E** Correlation between condition differences in modulation of SMA -> CB connection and learning index.

#### Modulation of SMA to cerebellar connections associated with better learning

We then asked whether condition differences in modulatory effects on connections from M1, SMA and PMC to CB reflect implicit learning of the underlying sequence. As no group differences were evident, we correlated the Learning Index (see Table 4) across all subjects of both groups, with the individual condition differences in modulatory parameters. We found a significant negative correlation in a connection from right SMA to left cerebellum (Fig. 4E, r = -0.46, p = 0.01) and a tendency for a negative correlation in a connection from left SMA to right cerebellum (Fig. 4D, r = -0.36, p = 0.05). These negative correlations suggest that the stronger the difference in modulation between SEQ and RND, the better the subject learnt.

In addition, we tested the hypothesis that connections from putamen (Pu) to thalamus (TH) are altered in PD compared to controls. There were no differences in left Pu -> left TH connection (controls 0.04 ± 0.03; PD: 0.02 ± 0.01, p = 0.4), as well as in right Pu -> right TH connection (controls 0.02 ± 0.02; PD: 0.05 ± 0.01, p = 0.3). In terms of the driving input to bilateral cerebellum by the task, we found a significant increase in the effect of task input during RND compared to SEQ blocks (right cerebellum: F_1,27_ = 24.7, p < 0.001; left cerebellum: F_1,27_ = 16.3, p < 0.001). There were no differences between the groups (p > 0.1).

## 4. Discussion

This study aimed to investigate the neural correlates underlying implicit motor sequence learning (MSL) deficits in PD. We found that neither healthy controls nor pwPD demonstrated significant sequence-specific learning across blocks of practice. Although online MSL has been shown to be relatively preserved depending on task complexity in healthy elderly subjects (see King et al., 2013), this suggests that implicitly learning a 12-item finger movement sequence is too difficult a task in this elder age-group. However, subjects of both groups made consistently more errors during random blocks compared to sequence blocks, suggesting that at least some chunks of the sequence were encoded. In addition, we found general task performance (stimulus-response association) deficits in PD, which could be related to nigral neurodegeneration and resulting reduction of dopaminergic neurotransmission. Analysis of fMRI data showed that healthy controls showed increased activity across blocks of task performance in left cerebellum crus II, insula and hippocampus as well as right premotor cortex, suggesting a link between task-related deficits and recruitment of these regions. In contrast, pwPD showed decreased activity in these regions. Indeed, we found that activity decrease over time in left hippocampus was associated with worse task performance over time in the PD group, highlighting a possible role of hippocampus in deficient learning of stimulus-response associations in PD. Importantly, activation of the substantia nigra (SN) during sequence learning was evident in the PD group only, and the magnitude of its activation was inversely correlated with learning. In addition, this increased activity in SN tended to correlate with LED and disease duration. One way to interpret this finding is that disease progression leads to “over-recruitment” of the remaining viable dopaminergic neurons in SN during sequence learning. However, whether degeneration of nigral dopaminergic neurons or over-recruitment of viable SN neurons is connected with learning deficits remains open. Finally, we found no differences between pwPD and controls in terms of effective connectivity in the cortico-striato-thalamo-cerebellar network. In both groups, the model with connections from M1 and premotor areas to cerebellum had the highest protected exceedance probability, suggesting that both pwPD and controls modulate the same cortico-cerebellar circuit during task performance. Modulation of connections from supplementary motor areas (SMA) to cerebellum across subjects of both groups was also correlated with the Learning Index (Table 4), such that better learners had larger differences in modulation due to sequence learning. This further supports the role of cortico-cerebellar connections in implicit MSL.

### Evidence for implicit motor sequence learning in PD and healthy controls

In general, reaction times (RT) in pwPD were significantly slower compared to healthy controls. General slowness probably reflects both bradykinesia and cognitive deficits in executive functions in PD (Cooper et al., 1994; Leis et al., 2005). On the group level, neither pwPD nor age-matched healthy controls improved RT during performance of the implicit 12-element sequence, when compared to random trials with no underlying sequential pattern. However, subjects of both groups made significantly more errors during random (RND) compared to sequence (SEQ) blocks. This effect is consistent with our previous findings in MSL tasks in healthy young subjects, in which larger error-rates in RND compared to SEQ blocks were found (Liebrand et al., 2020; Tzvi et al., 2016, 2015). Previously, we interpreted this effect as an implicit attempt to perform the sequence or chunks of the sequence in RND trials following sequence learning, however unsuccessfully. Thus, these results indicate that despite a lack of improvement in RT, some implicit sequence-specific learning was evident in all subjects. Note that using this exact same task, we have previously shown that younger healthy controls were faster during SEQ compared to RND blocks, while persons with cerebellar degeneration were not (Tzvi et al., 2017). Together with the current results, this suggests that age and neurological deficits impact on the ability to improve RT in this task. The absence of sequence learning differences between pwPD and their healthy controls contrasts with common findings showing task-specific learning deficits in PD using various sequence lengths, and mixtures of SEQ and RND blocks (for a systematic review see Ruitenberg et al., 2015). However, not all studies report the error-rates, which could also be an indicator for learning. For example, using a 12-element sequence of button presses and a mix of RND and SEQ blocks, Seidler and colleagues (2007) found that pwPD did not speed up performance in SEQ compared to RND blocks, while healthy controls did. Importantly, the authors found increased error-rates in RND compared to SEQ blocks in both groups, in line with the findings reported here.

In terms of general task performance (i.e., independent from condition), healthy controls showed significantly improved RT at the end of the task compared to the beginning, i.e. a better Performance Index (Table 4), while pwPD did not improve RT across blocks of task execution. In addition, a worse Performance Index in PD correlated (on trend level, p = 0.05) with their daily LED. As higher LED may be indicative of more advanced disease stages, this suggests that the ability to learn simple stimulus-response associations is affected by disease progression. Notably, this effect could not be explained by PD-associated reduced motor abilities (i.e., reduced movement speed due to bradykinesia), as the UPDRS-III score showed no correlation with the Performance Index.

### Decreased hippocampal activity is associated with deficient stimulus-response associations in PD

To obtain mechanistic insights into the possible stimulus-response deficits in PD, we analyzed differences in task-related changes of neural activity over time, between pwPD and healthy controls. We found large clusters in left cerebellum crus II, right premotor cortex, left hippocampus and left insula, along with other smaller clusters. Specifically, these regions showed increased activity in healthy subjects, and decreased activity in PD, suggesting that deficient learning of stimulus-response associations in PD is related to decreased activity in these regions. Indeed, we found that activity decrease over time in left hippocampus was associated with worse task performance over time in the PD group. The hippocampus is well-known for its important role in spatial memory encoding (Burgess et al., 2002), as well as in rapid acquisition of visuomotor mappings (Wise and Murray, 1999). Progressive hippocampal atrophy has been previously observed in pwPD (Brück et al., 2004; Camicioli et al., 2003) and was related to impaired memory (Jokinen et al., 2009). Calabresi et al. (2013) suggest that imbalanced interaction between dopamine transmission and hippocampal synaptic plasticity might play a role in learning and memory deficits observed in PD. Thus, our results may suggest that deficient ability to learn stimulus-response mappings is associated with hippocampal dysfunction in pwPD. Interestingly, task-related decrease in activity of left cerebellar crus II was significantly correlated with indirect markers of disease progression (LED, disease duration). This may indicate that intact endogenous nigro-striatal dopaminergic stimulation is necessary to adequately recruit cerebellar crus II across repeated task execution. This effect may be related to well described associations of pathological changes in the cerebellum in PD with dopaminergic degeneration, abnormal drives from the subthalamic nucleus, and dopaminergic treatment (see review by (Wu and Hallett, 2013)).

### Increased substantia nigra activity is predictive of motor sequence learning impairment in PD

Investigation of the underlying differences between pwPD and healthy controls in neural responses to MSL have revealed sequence-specific increased activity in a cluster with a peak in the right substantia nigra (SN) and adjacent bilateral midbrain areas in pwPD. Correlation analysis in this cluster showed that stronger activity in SN was associated with worse sequence-specific learning in the PD group. While activity increase in SN tended to correlate with LED and disease duration, no relationship between the Learning Index (Table 4) or the UPDRS-III score with LED was found. Daily dosages of dopaminergic medication as well as disease duration likely indicate the state of dopaminergic denervation in SN and thus the dysfunction of endogenous nigro-striatal dopaminergic stimulation. Previous studies show that lesions to SN in a rat model of PD lead to loss of striatal dopamine and deficits linked to habit learning and working memory (Da Cunha et al., 2002). In addition, recordings of SN neurons in humans performing a memory task showed that dopaminergic neurons in SN are modulated by memory (Kamiński et al., 2018). Matsumoto and colleagues (Matsumoto et al., 1999) showed that depletion of nigrostriatal dopamine in primates leads to a specific deficit in learning new motor sequences. Interestingly, while exogenous dopamine replacement therapy could restore high-speed walking after genetic ablation of dopaminergic neurons in the SN of mice, motor skill learning remained severely impaired (Wu et al., 2019). This may suggest that the process of acquiring new motor skills relies on specific learning-related dynamics of endogenous nigro-striatal dopamine release that cannot be compensated by exogenous dopamine replacement therapy (although cardinal motor symptoms of PD such as bradykinesia are effectively attenuated). In sum, the above body of evidence indicates that impairments of the capacity to learn sequential movements in PD is related to increasing demands on the dysfunctional endogenous dopaminergic system. Therefore, the correlation between LED/disease duration and activity increase in SN during sequence performance found in the current study may point to an increased (yet insufficient) drive to recruit the dysfunctional endogenous nigro-striatal dopaminergic system that is essential for MSL. As all pwPD were on their regular dopaminergic medication during the experiment, it seems improbable that increased SN activity and the associated impairment in MSL were a consequence of insufficient global dopamine availability during learning. Our finding rather supports the notion that MSL relies on sequence learning-specific recruitment of the endogenous nigro-striatal dopaminergic system.

### Implicit motor sequence learning modulates cortico-cerebellar connections independent from PD

We examined potential connectivity differences within the cortico-striato-thalamo-cerebellar network which might explain MSL deficits in PD. Specifically, we asked whether motor task performance modulated connections in the “basal-ganglia pathway”, in which the motor command propagates from motor and premotor areas to putamen, thalamus and back to cortical motor areas (Calabresi et al., 2014), or rather in the “cerebellar pathway”, in which motor commands are propagated from motor and premotor areas to cerebellum, and back through the thalamus (Ramnani, 2006). We expected to find evidence for abnormal striatal activity in PD reflected in connections with putamen. Using Bayesian model selection, we found that in the optimal model in both pwPD and controls connections from M1 and premotor areas to cerebellum were modulated by the motor task, replicating our previous findings (Liebrand et al., 2020; Tzvi et al., 2017, 2015, 2014). Importantly, we found no evidence for differences in modulatory parameters between PD and controls. It is plausible that unmodelled interactions from SN to putamen would have also revealed specific learning-related changes, but as the SN is too small to be modelled using DCM, it is not possible to examine this hypothesis using the data in this study. Nonetheless, the results of Bayesian model selection suggest that both pwPD and healthy controls implement the motor task using the same circuitry.

In addition, we found condition differences in both healthy controls and pwPD, in modulation of bilateral connections from M1, premotor cortex and SMA to cerebellum. Specifically, during random blocks negative modulation of these connections was evident, whereas modulation by sequence blocks was close to zero. These results are in accordance with our previous findings in persons with cerebellar degeneration (Tzvi et al., 2017) but in contrast to our findings in healthy young controls (Tzvi et al., 2014), in which negative modulation of this connection was observed during sequence blocks. Together with the lack of RT differences between SEQ and RND blocks, these results suggest that previously found negative modulation in SEQ relate to improved RT, whereas here, negative modulation during RND could be related to higher error-rates. Importantly, we observed that modulation of SMA to cerebellum connection was associated with the Learning Index (Table 4), such that subjects who were better in sequence learning, also had stronger condition differences in modulation of this connection, probably driven by larger negative modulation during RND. Given the important role of SMA in forming sequential representations of movements (Cona and Semenza, 2017), it is conceivable that subjects who better encoded the underlying sequence also implicitly expected sequential patterns to appear during RND. The lack of sequential pattern then led to decreased communication from SMA to cerebellum.

### Limitations

Some limitations of this study need to be mentioned. First, the lack of RT differences between SEQ and RND in both pwPD and healthy controls suggest that on the group level no learning has taken place. Note however, that error-rate differences between conditions can also be indicative of learning, albeit indirectly. Second, although 23 pwPD were originally included in the study, for various reasons, the data of only 16 pwPD could be analyzed. This relatively small sample size may have led to the marginal differences between pwPD and controls in terms of neural activity underlying general task improvements, as well as the trending but non-significant relationships with parameters of disease progressions (LED, disease duration). Third, we examined several relationships between behavioral data as well as activation changes and parameters of disease progressions. As these tests were exploratory, we did not perform any corrections for multiple comparisons. Future studies should scrutinize the relationship between behavioral measures of MSL, changes in activity and parameters of disease progressions.

## Conclusions

In this study we investigated the neural correlates underlying implicit MSL deficits in PD. Increase in error-rates during RND compared to SEQ in both groups suggested that some sequence learning was evident. FMRI analysis revealed two noteworthy findings: first, pwPD showed decreased hippocampal activity with time, and this decrease was associated with worse task performance. Second, pwPD exhibited abnormal activity increase in SN during SEQ blocks specifically, which was associated with worse learning of the underlying sequence. This sequence-specific increase in SN activity in pwPD may point to an increased drive to recruit the dysfunctional endogenous nigro-striatal dopaminergic system for the purpose of learning. Finally, despite behavioral and activation differences between PD and controls, we found no group differences in connectivity within a cortico-striato-thalamo-cerebellar network. Importantly, the same model was evident in both groups suggesting that despite the dysfunctional dopaminergic system, pwPD may implement the same circuitry for motor task performance.

## Abbreviations

DCM: dynamic causal modelling
fMRI: functional magnetic resonance imaging
LED: levodopa equivalent dose
MSL: motor sequence learning
pwPD: persons with Parkinson’s disease
RT: reaction times
RND: random
SEQ: sequence
SN: substantia nigra;

## Acknowledgements

We’d like to thank all participants for their time. We also thank Christoph Zimmermann, Sinem Tunc, Henrike Hanssen, Jannik Prasuhn, and Mourad Zoubir for their assistance with subject recruitment and data collection. This study was supported by internal funding of the University of Lübeck [grant J29-2017] and by the DFG [grant TZ 85/1-1] to ET. NB received speaker’s honoraria from Abbvie and Teva. He served as a consultant for Censa Pharmaceuticals and Centogene.

